# Thalamic Nuclei Functional Controllability Accounts for Cognitive Impairment in Multiple Sclerosis Over and Above Structural Damage

**DOI:** 10.64898/2026.05.01.722249

**Authors:** Yuping Yang, Anna Woollams, Ilona Lipp, Jessica Haigh, Rose-Marie Kouwenhoven, Valentina Tomassini, Nelson J Trujillo-Barreto, Nils Muhlert

**Author notes:** Senior author (Nils Muhlert). **Correspondence to:** Dr Yuping Yang, Division of Psychology, Communication and Human Neurosciences, Faculty of Biology, Medicine and Health, University of Manchester, Oxford Road, Manchester M13 9PL, UK, Dr Nils Muhlert, Division of Psychology, Communication and Human Neurosciences, Faculty of Biology, Medicine and Health, University of Manchester, Oxford Road, Manchester M13 9PL, UK.

## Abstract

**Background:** The thalamus has emerged as a key region involved in cognitive dysfunction in multiple sclerosis (MS). While previous studies have identified associations between thalamic structural damage, altered functional connectivity, and cognitive performance, the specific contributions of individual thalamic nuclei and the added value of integrating structural and functional metrics remain poorly understood.

**Methods:** T1-weighted MRI, diffusion MRI, resting-state fMRI, and neuropsychological data were collected from 102 individuals with MS and 27 healthy controls. Thalamic grey matter volume, white matter microstructural integrity, and functional controllability were calculated for each nucleus and compared between individuals with MS and healthy controls, as well as between MS cognitive subgroups. Partial Spearman correlations were used to examine the relationship between imaging metrics across the three modalities, and also between imaging metrics and cognitive performance in MS. Sparse canonical correlation analysis models were used to examine the covariance between thalamic imaging metrics and cognitive performance in MS.

**Results:** Widespread atrophy and microstructural damage were observed across all thalamic nuclei in individuals with MS, regardless of cognitive status. In contrast, alterations in functional controllability were more spatially specific, primarily affecting the medial dorsal anterior nuclei, and were most pronounced in cognitively impaired individuals. These functional controllability metrics were independent of grey matter volume, white matter integrity, and lesion load. Combining thalamic functional controllability with structural metrics yielded a stronger association with cognitive performance in MS than either modality alone.

**Conclusion:** This study provides novel evidence that functional controllability in the thalamus, particularly within the medial dorsal anterior nuclei, plays a critical role in cognitive impairment in MS. By applying a network control framework, our findings offer a dynamic systems perspective that extends beyond traditional connectivity analyses, capturing the thalamus’s role in supporting flexible cognitive transitions. The integration of structural and functional controllability metrics enhances the ability to characterise individual differences in cognitive performance and may inform future efforts to identify biomarkers of cognitive dysfunction in MS.

## Introduction

Multiple sclerosis (MS) is a progressive inflammatory and degenerative disease characterised by demyelination and axonal loss in both brain white and grey matter (Fillippi et al., 2018). Among the deep brain structures affected in MS, the thalamus has received particular attention, as it is commonly involved from the earliest stages of the disease (Azevedo et al., 2018; Coupé et al., 2023; Eshaghi et al., 2018; Tozlu et al., 2023) and across its various subtypes (Azevedo et al., 2018; Eshaghi et al., 2018; Sepulcre et al., 2006).

Cognitive impairment is a frequent symptom of MS, affecting between 40% and 70% of individuals (Rocca et al., 2015). This has led to extensive investigation into the neural mechanisms underlying cognitive dysfunction in MS, with mounting evidence pointing to the thalamus as a key contributor (reviewed in (Amin & Ontaneda, 2021; DeLuca et al., 2015; Kipp et al., 2015)). As a central hub for information modulation across the brain, the thalamus plays a crucial role in human cognition, with its multiple nuclei supporting various cognitive processing such as memory, attention, executive function, language processing, and visuospatial function (reviewed in (Shine et al., 2023; Wolff & Vann, 2019)). Damage to the thalamus has been consistently linked to poorer performance across these domains in MS, highlighting its important role in cognitive impairment in the disease (Bergsland et al., 2021; Bisecco et al., 2021; Jandric et al., 2022; Lin et al., 2019; Lorefice et al., 2020; Schoonheim et al., 2015; Yang et al., 2025).

Thalamic structural damage is one of the most consistently reported indicators of cognitive impairment in MS. Numerous structural MRI studies have identified thalamic volume atrophy (Bergsland et al., 2016, 2021; Bisecco et al., 2021; Houtchens et al., 2007; Lorefice et al., 2020; Rocca et al., 2010; Rojas et al., 2018; Schoonheim et al., 2015; Štecková et al., 2014), thalamic fractional anisotropy (FA) and mean diffusivity (MD) changes (Bergsland et al., 2018; Conway et al., 2021; Schoonheim et al., 2014) as strong factors that associated with cognitive performance in people with MS. Thalamic functional alterations have also received growing attention in MS research over the past two decades. Recent functional MRI studies have shown that increased connectivity between the thalamus and the inferior frontal gyrus is associated with poorer performance in executive functioning, information processing speed, visuospatial memory, working memory, and psychomotor speed (Schoonheim et al., 2015), whereas decreased functional connectivity between these regions has been linked to better performance in visual memory and global cognition (Rocca et al., 2018). Besides, increased functional thalamocortical connectivity has been associated with poorer auditory information processing speed (Tona et al., 2014). Additionally, increased functional connectivity between the thalamus and the right frontal/postcentral cortex (Carotenuto et al., 2022), as well as between the thalamus and the superior temporal cortex (d’Ambrosio et al., 2017) has been observed only in cognitively impaired (CIMS) but not cognitively preserved individuals with MS (CPMS). However, these functional connectivity analyses primarily reflect synchronous activity during rest and therefore fail to capture the thalamus’s role in modulating dynamic control and transitions between cognitive states, which are crucial for the efficiency and flexibility of cognitive performance.

Network controllability analysis has emerged as a promising approach for characterising how specific regions influence the brain dynamic control and transitions between cognitive states (Lynn & Bassett, 2019; Gu et al., 2015; Deng et al., 2022). This method has been shown powerful to explain cognitive performance in healthy individuals (Deng et al., 2022; Deng & Gu, 2020; Gu et al., 2015)(Yang et al., 2025. bioRxiv) and has demonstrated promise in identifying brain biomarkers across various clinical conditions (Bernhardt et al., 2019; Hahn et al., 2023; Li et al., 2023; Parkes et al., 2021; Zarkali et al., 2020). In MS, recent evidence suggests that cognitively impaired individuals experience greater difficulty transitioning between brain states compared to cognitively preserved individuals (Broeders et al., 2024), with the thalamus identified as a key region where altered controllability may constrain the brain’s capacity to perform cognitively demanding, high-energy-cost transitions (Yang et al., 2025).

Current evidence collectively highlights the importance of the thalamus as a whole, in terms of both structure and function, in MS-related cognitive impairment; however, the specific contributions of individual thalamic nuclei, as well as the added value of combining their structural and functional properties, remain poorly understood. Combining structural and functional metrics of thalamic nuclei has shown promise in elucidating cognitive performance in healthy individuals (Yang et al., 2025, bioRxiv. Chapter 5) and in MS (Schoonheim et al., 2015). Building on this, the present study integrates three neuroimaging modalities—grey matter volume, diffusion-based white matter integrity, and functional controllability—to investigate MS-related alterations within individual thalamic nuclei. We then examine how these metrics, both independently and in combination, relate to inter-individual variability in cognitive performance. We hypothesise that (1) thalamic grey matter volume, white matter microstructural integrity, and functional controllability in specific nuclei are altered in individuals with MS compared to healthy controls, with more pronounced alterations in functional controllability metrics among those with cognitive impairment; (2) thalamic functional controllability metrics captures unique aspects of MS-related thalamic pathology that are not accounted for by grey or white matter structural changes or lesion load; and (3) combining thalamic functional controllability metrics and structural grey and white matter metrics yields a stronger association with cognitive performance over either modality alone.

## Materials and Methods

### Participants

A total of 102 patients with relapsing-remitting MS were recruited from the Helen Durham Centre for Neuroinflammation at the University Hospital of Wales, alongside 27 healthy controls (HC) recruited from the local community. All participants were right-handed, aged between 18 and 60 years, and had no contraindications to MRI. Additional inclusion criteria for patients included no comorbid neurological or psychiatric conditions, no changes to disease-modifying treatments within three months prior to scanning, and a relapse-free period at the time of participation. All participants underwent demographic, clinical, and psychological assessments, as well as MRI scanning, during a single study visit.

The study received ethical approval from the NHS South-West Research Ethics Committee and the Cardiff and Vale University Health Board R&D Committee. Written informed consent was obtained from all participants.

### Behavioural data

Demographic variables included age, sex, and years of education. Clinical measures comprised disease duration, Expanded Disability Status Scale (EDSS) scores, and Multiple Sclerosis Functional Composite (MSFC) scores. Cognitive performance was assessed using the Brief Repeatable Battery of Neuropsychological Tests (BRB-N), which includes nine subtests covering four cognitive domains: (1) verbal memory, assessed by the Selective

Reminding Tests; (2) visual memory, evaluated using the 10/36 Spatial Recall Tests; (3) attention, information processing, and executive function, measured by the Symbol Digit Modalities Test and Paced Auditory Serial Addition Tests; and (4) verbal fluency, assessed via the Word List Generation Test. To evaluate cognitive impairment in patients, raw scores on each BRB-N subtest were converted into Z-scores using the mean and standard deviation (SD) of the healthy control group. Patients were classified as cognitively impaired multiple sclerosis (CIMS) if they scored ≥1.5 SDs below the control mean on at least two subtests; all other patients were categorised as cognitively preserved MS (CPMS).

### MRI data acquisition

MRI data were acquired using a 3T General Electric HDx MRI system (GE Medical Systems, Milwaukee, WI) equipped with an 8-channel receive-only head radiofrequency coil. A high-resolution 3D T1-weighted sequence (3DT1) was acquired for lesion identification, tissue segmentation, and spatial registration (voxel size = 1 mm × 1 mm × 1 mm; echo time [TE] = 3.0 ms; repetition time [TR] = 7.8 ms; matrix = 256 × 256 × 172; field of view [FOV] = 256 mm × 256 mm; flip angle [FA] = 20°).

A T2/proton density–weighted sequence (voxel size = 0.94 mm × 0.94 mm × 4.5 mm, TE = 9.0/80.6 ms, TR = 3,000 ms, FOV = 240 mm × 240 mm, number of slices = 36, FA = 90°) and a fluid-attenuated inversion recovery (FLAIR) sequence (voxel size = 0.86 mm × 0.86 mm × 4.5 mm, TE = 122.3 ms, TR = 9,502 ms, FOV = 220 mm × 220 mm, number of slices = 36, FA = 90°) were acquired for identification and segmentation of T2-hyperintense MS lesions.

Resting-state functional MRI (rs-fMRI) was performed using a T2*-weighted gradient-echo echo-planar imaging sequence (voxel size = 3.4 mm × 3.4 mm × 3 mm, TE = 35 ms, TR = 3,000 ms, matrix size = 64 × 64 × 46, FOV = 220 mm × 220 mm, number of volumes = 100, number of slices = 46, interleaved order). Participants were instructed to remain relax with their eyes closed during rs-fMRI acquisition.

Diffusion-weighted MRI (dMRI) was acquired using a twice-refocused spin-echo echo-planar imaging sequence with 6 volumes without diffusion weighting (b = 0 s/mm²) and 40 volumes with diffusion gradients applied in uniformly distributed directions (b = 1,200 s/mm²; voxel size = 1.8 mm × 1.8 mm × 2.4 mm; TE = 94.5 ms; TR = 16,000 ms; FOV = 230 mm × 230 mm; number of slices = 57).

### MRI data preprocessing

Structural 3DT1 data from patients were lesion-filled using the method described previously(Lipp et al., 2019) to improve the accuracy of subsequent tissue segmentation.

Lesion-filled images were then segmented into grey matter (GM), white matter (WM), and cerebrospinal fluid (CSF) using FSL’s Automated Segmentation Tool (FAST). Segmentation quality was assessed manually for all participants. Binary masks of intracranial tissue, excluding CSF, were generated by combining the GM and WM segmentations and used for downstream diffusion MRI analyses. Lesion volumes were calculated from the binary lesion masks generated during the lesion-filling process. A final spatial smoothing step was applied using a Gaussian kernel with a full width at half maximum (FWHM) of 6 mm.

Preprocessing of dMRI data was performed using ExploreDTI as described in our previous work.(Jandric et al., 2021) The pipeline included correction for head motion, eddy current distortions, and echo planar imaging (EPI)–induced geometric distortions. Each diffusion-weighted image was registered to its corresponding skull-stripped and downsampled (to 1.5 mm isotropic resolution) 3D T1-weighted image using Elastix. Diffusion-encoding vectors were reoriented accordingly to preserve anatomical accuracy following spatial transformations. All processed dMRI data underwent manual quality control to ensure data integrity and exclude artefacts.(Jandric et al., 2021)

rs-fMRI data were preprocessed using SPM1, as described in our previous work.(Yang et al., 2025) Briefly, functional images were corrected for head motion and slice timing differences. There were no significant group differences in either maximum or mean frame-wise displacement (p > 0.05, permutation test). Motion-corrected images were spatially normalised to the MNI152 space using deformation fields derived from tissue segmentation of the corresponding structural T1-weighted images. The normalised functional images were then spatially smoothed using a Gaussian kernel with a 6-mm FWHM.

### Thalamic volume

Following preprocessing, brain volume maps were generated from lesion-filled 3DT1 images using FSL’s SIENAX tool. Subcortical structures were then parcellated into 16 thalamic nuclei, eight per hemisphere, based on a standard subcortical parcellation atlas,(Tian et al., 2020) as previously described.(Yang et al., 2025) The identified nuclei included (1) two nucleus per hemisphere in the dorsoanterior thalamus, namely medial dorsoanterior thalamus (THA-DAm-rh/ THA-DAm-lh) and lateral dorsoanterior thalamus (THA-DAl-rh/ THA-DAl-lh); (2) three nucleus per hemisphere in the ventroanterior thalamus, namely superior ventroanterior thalamus (THA-VAs-rh/ THA-VAs-lh), anterior inferior ventroanterior thalamus (THA-VAia-rh/ THA-Vaia-lh), and posterior inferior ventroanterior thalamus (THA-VAip-rh/ THA-VAip-lh); (3) one nuclei per hemisphere in the dorsoposterior thalamus (THA-DP-rh/ THA-DP-lh); and (4) two nucleus per hemisphere in the ventroposterior thalamus, namely medial ventroposterior thalamus (THA-VPm-rh/ THA-VPm-lh) and lateral ventroposterior thalamus (THA-VPl-rh/ THA-VPl-lh). For each nucleus, volume was computed by averaging the voxel-wise grey matter volume within its defined anatomical boundaries.

### Thalamic structural integrity

Diffusion tensors were fitted using FSL’s *DTIFIT* to generate fractional anisotropy (FA) and mean diffusivity (MD) maps. FA and MD values specific to each thalamic nucleus were extracted by applying the corresponding track masks derived from tractography. Whole-brain FA and MD maps served as inputs to FSL’s *BedpostX* pipeline, which models white matter fibre orientations and crossing fibres for probabilistic tractography. The resulting outputs were subsequently used in FSL’s *ProbtrackX2* for tractography, as described previously.(Yang et al., 2025) For each thalamic nucleus, tractography was performed using the nucleus as a seed mask, with waypoint masks identified via the Functionnectome toolbox.(Nozais et al., 2023) The cerebellum, brainstem, and ventricles were defined as termination masks and excluded from further analysis to restrict streamlines to cortical and subcortical regions, consistent with procedures applied in the volume and functional analyses. Spatial linear and non-linear transformations between standard, diffusion, and individual anatomical spaces were computed using *FLIRT* and *FNIRT* and applied throughout the tractography process. For each voxel within a seed region, 5,000 streamlines were initiated with a step length of 0.5 mm and a maximum of 2,000 steps per streamline.

Streamline propagation was constrained by a curvature threshold of 0.2 and a minimum fibre orientation volume fraction threshold of 0.01. This process yielded probabilistic streamline maps for each thalamic nucleus. Track density (TD) was quantified by averaging the probabilistic streamlines passing through the respective nucleus.

### Thalamic functional controllability

To investigate the controllability of individual thalamic nuclei over brain dynamics, a functional connectivity network was constructed for each participant using a combined parcellation of 454 regions of interest (ROIs). This comprised 400 cortical ROIs defined by the Schaefer atlas, which captures canonical functional organization(Schaefer et al., 2018), and 54 subcortical ROIs based on a connectivity-gradient parcellation.(Tian et al., 2020) Voxel-wise BOLD signals were averaged within each ROI, and pairwise Pearson correlation coefficients were calculated to define the functional connectivity matrix. Using a simplified linear time-invariant control model(Deng et al., 2022; Gu et al., 2015), controllability metrics for each thalamic nucleus were estimated to characterize their influence on brain activity transitions. Three metrics were derived: average controllability, modal controllability, and activation energy, as detailed in our previous work.(Yang et al., 2025) Average controllability quantifies a node’s capacity to drive the system toward easily reachable states with low energy input, whereas modal controllability reflects the ability to steer the system toward difficult-to-reach states requiring higher energy. Activation energy represents the minimal energy required to activate a node to control system dynamics (see Fig. 1 and Supplementary Methods for further details).(Deng et al., 2022; Gu et al., 2015)

**^1^Figure 1.**
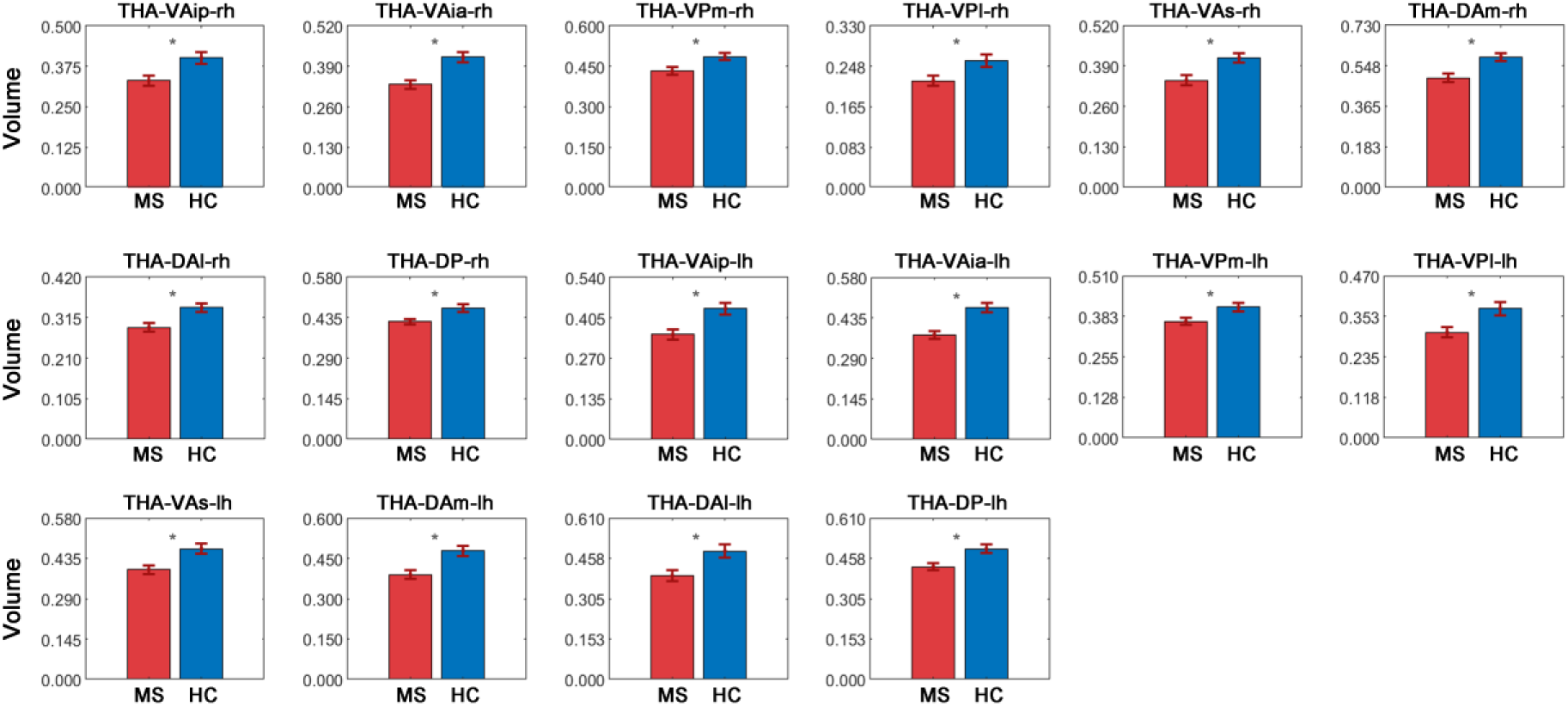
Thalamic volume changes in MS. Significant decreases in volume were observed in MS in all thalamic nuclei.

### Statistical analysis

All statistical procedures were carried out in MATLAB R2022b (MathWorks, Inc.). Chi-square tests were employed to compare categorical variables (e.g., sex). Permutation testing with 10,000 repetitions assessed group differences in continuous demographic, clinical, and neuropsychological variables. Neuropsychological analyses accounted for age and sex as covariates. Results were presented with 95% confidence intervals.

#### Thalamic imaging metrics changes in MS

Group differences in thalamic imaging metrics (volume metrics; diffusion metrics: FA, MD, TD; and functional controllability metrics: average controllability, modal controllability, activation energy) between MS and HC were assessed using non-parametric permutation t-test (10,000 times). Initially, two-sample t-tests were conducted for each imaging metric to obtain observed group differences. To estimate the empirical null distribution, group labels were randomly shuffled, and the t-statistic recomputed across 10,000 permutations. The empirical *p*-value was defined as the proportion of permuted t-values that were equal to or greater than the absolute value of the original test statistic. Age and sex were included as covariates in the analysis. Multiple comparisons were corrected using the Bonferroni method, applying a significance threshold of *p* < 0.05.

#### Thalamic imaging metrics changes in CIMS and CPMS

To explore how thalamic imaging metrics changes relate to cognitive impairment in MS, we further assessed thalamic imaging metrics difference between CIMS, CPMS and HC. Specifically, non-parametric permutation f-tests (10,000 times) were first applied between CIMS, CPMS and HC, following the procedures described above. For metrics showing significant group effects after Bonferroni correction, post hoc pairwise comparisons were carried out using permutation t-test (10,000 times). Again, age and sex were controlled for as covariates in all analyses.

#### Associations between thalamic imaging metrics and clinical/cognitive outputs

To investigate the relationship between thalamic imaging metrics and clinical/cognitive outputs in MS, correlation analyses were performed between imaging metrics (volume metrics; diffusion metrics: FA, MD, TD; and functional controllability metrics: average controllability, modal controllability, activation energy) and clinical variables (Lesion load, disease duration, EDSS, MSFC) or cognitive performance of each of the domains (Verbal memory, Visual memory, AtteExec, and VerbFlue), in MS group and CIMS group. Variable distributions were first assessed using the Lilliefors test. For variables that met the assumption of normality, Pearson correlation coefficients were computed; otherwise, Spearman correlations were used. All analyses involving neuropsychological and imaging metrics were adjusted for age and sex.

#### Associations among thalamic imaging metrics

To explore whether metrics derived from different imaging modalities captured overlapping or distinct aspects of the thalamic nuclei relevant to cognition, correlation analyses were also conducted among thalamic volume metrics, diffusion metrics (FA, MD, TD), and functional controllability metrics (average controllability, modal controllability, activation energy) in MS group and CIMS group. Normality was assessed using the Lilliefors test, with Pearson or Spearman correlations applied accordingly. Age and sex were included as covariates in all imaging-related analyses.

#### Covariance between thalamic imaging metrics and cognitive performance

Sparse canonical correlation analysis (CCA) (Mackey, 2008; Uurtio et al., 2019) was employed to assess the multivariate associations between imaging metrics and cognitive domains in MS. Specifically, we assessed whether single-modality or multi-modality models better capture the covariance with cognitive function. Imaging inputs were grouped as: 1)

Single modality: volume metrics only, diffusion metrics (FA, MD, TD) only, or functional controllability metrics (average controllability, modal controllability, activation energy) only; 2) Dual modality: volume + diffusion, volume + functional, or diffusion + functional; and 3) Triple modality: volume, diffusion, and functional metrics combined. Prior to analysis, all variables were transformed into standard scores (z-scores). Sparse CCA was employed to uncover multivariate associations between imaging metrics and cognitive performance, using an alternating projected gradient method. Regularisation was optimised via grid search combined with 10-fold cross-validation. Initialisation involved both singular vectors and randomly generated vectors derived from the cross-covariance matrix. Model selection was based on the average test-set correlation across cross-validation folds (Mackey, 2008; Uurtio et al., 2019).

## Results

### Thalamic imaging metrics changes in MS

#### Thalamic volume metrics changes in MS

Significant decreases in volume were observed in MS in all thalamic nuclei (Figure 1): THA-VAip-rh (t = -4.069, p < 0.001), THA-VAia-rh (t = -5.704, p < 0.001), THA-VPm-rh (t = -3.591, p < 0.001), THA-VPl-rh (t = -3.590, p < 0.001), THA-VAs-rh (t = -3.967, p < 0.001), THA-DAm-rh (t = -4.555, p < 0.001), THA-DAl-rh (t = -3.950, p < 0.001), THA-DP-rh (t = -4.120, p < 0.001), THA-VAip-lh (t = -4.909, p < 0.001), THA-VAia-lh (t = -5.991, p < 0.001), THA-VPm-lh (t = -3.931, p < 0.001), THA-VPl-lh (t = - 4.159, p < 0.001), THA-VAs-lh (t = -4.401, p < 0.001), THA-DAm-lh (t = -4.941, p < 0.001), THA-DAl-lh (t = -4.323, p < 0.001), and THA-DP-lh (t = -4.833, p < 0.001).

#### Thalamic diffusion metrics changes in MS

Significant decreases in FA were observed in MS in all thalamic nuclei (Figure 2A): THA-VAip-rh (t = -3.786, p < 0.001), THA-VAia-rh (t = - 3.801, p < 0.001), THA-VPm-rh (t = -3.321, p = 0.002), THA-VPl-rh (t = -3.137, p = 0.002), THA-VAs-rh (t = -3.664, p < 0.001), THA-DAm-rh (t = -3.552, p < 0.001), THA-DAl-rh (t = - 3.093, p = 0.002), THA-DP-rh (t = -3.420, p = 0.001), THA-VAip-lh (t = -3.517, p < 0.001), THA-VAia-lh (t = -3.351, p = 0.001), THA-VPm-lh (t = -3.243, p = 0.001), THA-VPl-lh (t = -2.971, p = 0.003), THA-VAs-lh (t = -3.309, p < 0.001), THA-DAm-lh (t = -3.358, p = 0.001), THA-DAl-lh (t = -3.366, p < 0.001), and THA-DP-lh (t = -3.496, p < 0.001). Besides, significant increases in MD were observed in MS in THA-VAia-lh (t = 2.873, p = 0.003) (Figure 2B). Additionally, significant changes in TD were observed in MS in THA-VPl-rh (t = 4.341, p < 0.001), THA-VPl-lh (t = 3.844, p < 0.001), and THA-DAm-lh (t = -3.115, p = 0.003) (Figure 2C).

**Figure 2.**
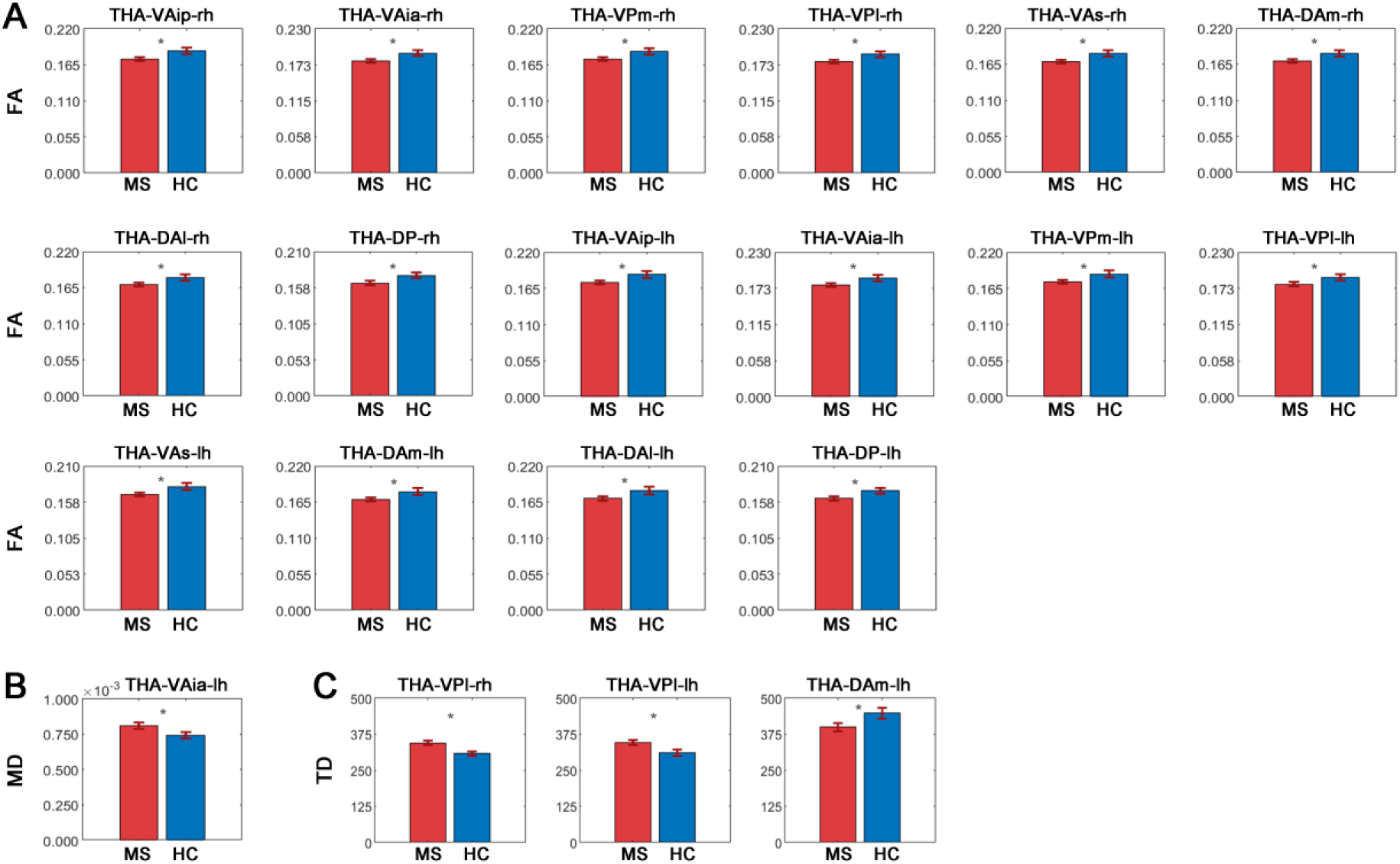
Thalamic diffusion metrics changes in MS. (2A) Significant decreases in FA were observed in MS in all thalamic nuclei. **(2B)** Significant increases in MD were observed in MS in THA-VAia-lh. **(2C)** Significant changes in TD were observed in MS in THA-VPl-rh, THA-VPl-lh, and THA-DAm-lh.

#### Thalamic controllability metrics changes in MS

Significant increases in average controllability were observed in MS in THA-DAm-rh (t = 3.345, p < 0.001) and THA-DAl-rh (t = 3.430, p < 0.001) (Figure 3A). Besides, significant decreases in modal controllability were observed in MS in THA-DAm-rh (t = -3.689, p < 0.001) and THA-DAl-rh (t = -3.224, p = 0.001) (Figure 3B). Additionally, significant decreases in activation energy were observed in MS in THA-DAm-rh (t = -3.479, p < 0.001) and THA-DAl-rh (t = -3.839, p < 0.001) (Figure 3C).

**Figure 3.**
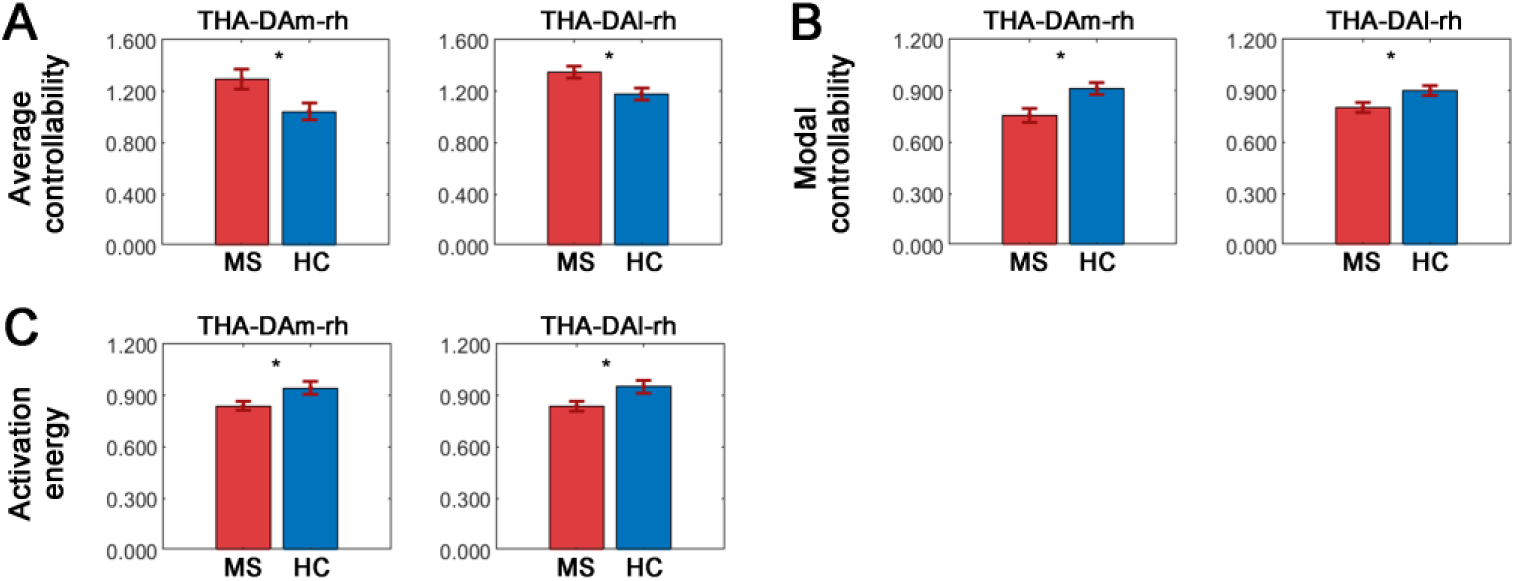
Thalamic controllability changes in MS. (3A) Significant increases in average controllability were observed in MS in THA-DAm-rh and THA-DAl-rh. **(3B)** Significant decreases in modal controllability were observed in MS in THA-DAm-rh and THA-DAl-rh. **(3C)** Significant decreases in activation energy were observed in MS in THA-DAm-rh and THA-DAl-rh.

### Thalamic imaging metrics changes in CIMS and CPMS

#### Thalamic volume metrics changes in CIMS and CPMS

Both the CIMS and CPMS groups exhibited significant volume decreases compared to HC in all thalamic nuclei, but no differences were found between CIMS and CPMS in any of the thalamic nuclei (Figure 4): THA-VAip-rh (CIMS vs HC: t = -3.843, p < 0.001; CPMS vs HC: t = -3.423, p < 0.001; CIMS vs CPMS: t = -0.421, p > 0.05), THA-VAia-rh (CIMS vs HC: t = -5.699, p < 0.001; CPMS vs HC: t = - 4.708, p < 0.001; CIMS vs CPMS: t = -1.030, p > 0.05), THA-VPm-rh (CIMS vs HC: t = -3.380, p = 0.001; CPMS vs HC: t = -3.152, p = 0.002; CIMS vs CPMS: t = -0.214, p > 0.05), THA-VPl-rh (CIMS vs HC: t = -3.395, p < 0.001; CPMS vs HC: t = -3.105, p = 0.003; CIMS vs CPMS: t = - 0.289, p > 0.05), THA-VAs-rh (CIMS vs HC: t = -3.708, p < 0.001; CPMS vs HC: t = -3.219, p = 0.001; CIMS vs CPMS: t = -0.472, p > 0.05), THA-DAm-rh (CIMS vs HC: t = -4.164, p < 0.001; CPMS vs HC: t = -4.115, p < 0.001; CIMS vs CPMS: t = -0.025, p > 0.05), THA-DAl-rh (CIMS vs HC: t = -3.574, p < 0.001; CPMS vs HC: t = -3.607, p < 0.001; CIMS vs CPMS: t = -0.091, p > 0.05), THA-DP-rh (CIMS vs HC: t = -3.809, p < 0.001; CPMS vs HC: t = -3.664, p < 0.001; CIMS vs CPMS: t = -0.116, p > 0.05), THA-VAip-lh (CIMS vs HC: t = -4.791, p < 0.001; CPMS vs HC: t = -4.105, p < 0.001; CIMS vs CPMS: t = -0.745, p > 0.05), THA-VAia-lh (CIMS vs HC: t = -5.906, p < 0.001; CPMS vs HC: t = -4.970, p < 0.001; CIMS vs CPMS: t = -1.027, p > 0.05), THA-VPm-lh (CIMS vs HC: t = -3.709, p < 0.001; CPMS vs HC: t = -3.444, p < 0.001; CIMS vs CPMS: t = - 0.253, p > 0.05), THA-VPl-lh (CIMS vs HC: t = -4.120, p < 0.001; CPMS vs HC: t = -3.894, p < 0.001; CIMS vs CPMS: t = -0.164, p > 0.05), THA-VAs-lh (CIMS vs HC: t = -4.399, p < 0.001; CPMS vs HC: t = -3.336, p = 0.002; CIMS vs CPMS: t = -1.161, p > 0.05), THA-DAm-lh (CIMS vs HC: t = -4.893, p < 0.001; CPMS vs HC: t = -4.113, p < 0.001; CIMS vs CPMS: t = -0.821, p > 0.05), THA-DAl-lh (CIMS vs HC: t = -4.170, p < 0.001; CPMS vs HC: t = -3.752, p < 0.001; CIMS vs CPMS: t = -0.420, p > 0.05), and THA-DP-lh (CIMS vs HC: t = -4.563, p < 0.001; CPMS vs HC: t = -4.078, p < 0.001; CIMS vs CPMS: t = -0.500, p > 0.05).

**Figure 4.**
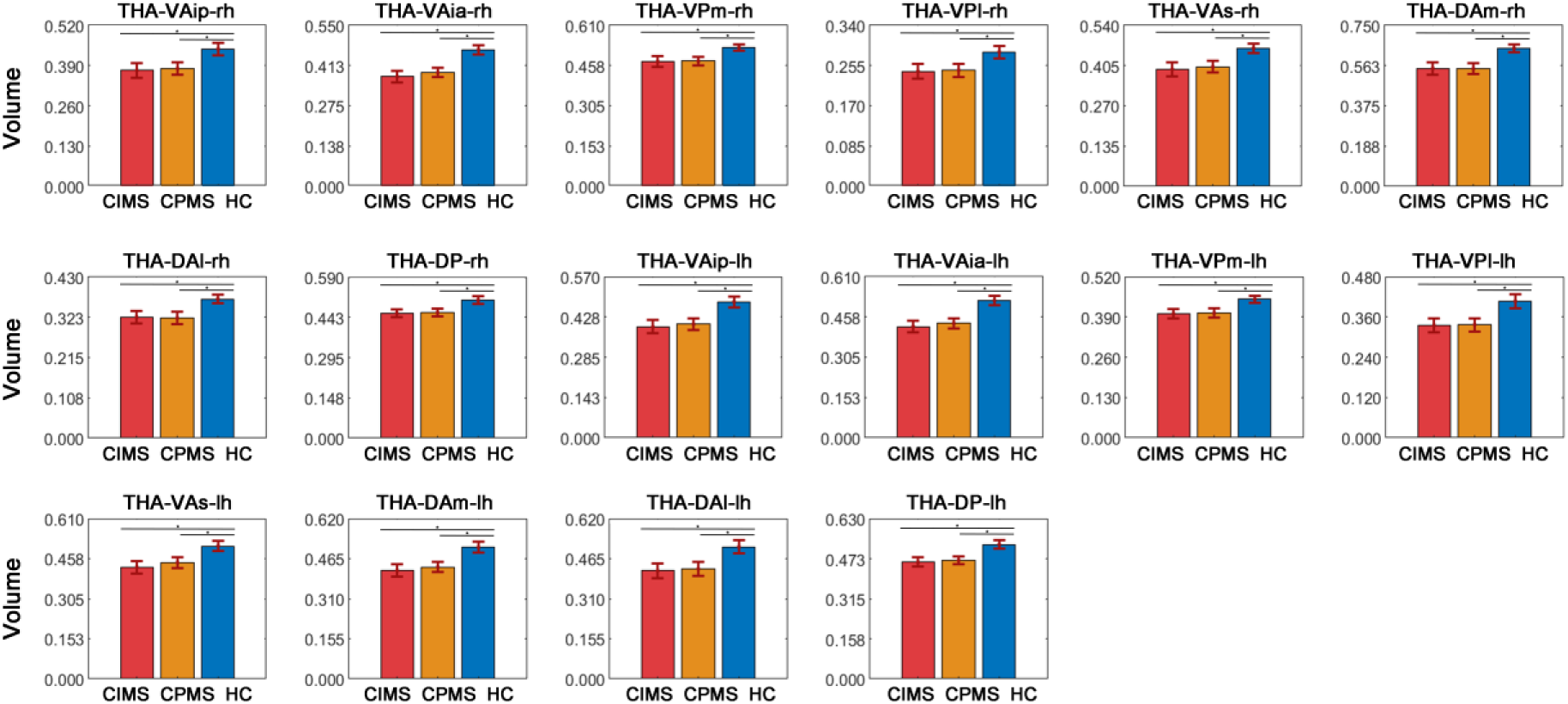
Thalamic volume changes in CIMS and CPMS. Both the CIMS and CPMS groups exhibited significant volume decreases compared to HC in all thalamic nuclei, but no differences were found between CIMS and CPMS in any of the thalamic nuclei.s

#### Thalamic diffusion metrics changes in CIMS and CPMS

Both the CIMS and CPMS groups exhibited significant FA decreases compared to HC in THA-VAip-rh, THA-VAia-rh, THA-VAs-rh, THA-DAm-rh, THA-DP-rh, THA-VAip-lh, and THA-DP-lh, but no differences were found between CIMS and CPMS in any of the thalamic nuclei (Figure 5A): THA-VAip-rh (CIMS vs HC: t = -3.735, p < 0.001; CPMS vs HC: t = -3.122, p = 0.002; CIMS vs CPMS: t = -0.676, p > 0.05), THA-VAia-rh (CIMS vs HC: t = -3.849, p < 0.001; CPMS vs HC: t = -3.036, p = 0.003; CIMS vs CPMS: t = -0.916, p > 0.05), THA-VAs-rh (CIMS vs HC: t = -3.637, p < 0.001; CPMS vs HC: t = - 2.996, p = 0.003; CIMS vs CPMS: t = -0.711, p > 0.05), THA-DAm-rh (CIMS vs HC: t = -3.481, p < 0.001; CPMS vs HC: t = -2.953, p = 0.004; CIMS vs CPMS: t = -0.578, p > 0.05), THA-DP-rh (CIMS vs HC: t = -3.362, p < 0.001; CPMS vs HC: t = -2.839, p = 0.006; CIMS vs CPMS: t = - 0.579, p > 0.05), THA-VAip-lh (CIMS vs HC: t = -3.482, p < 0.001; CPMS vs HC: t = -2.892, p = 0.004; CIMS vs CPMS: t = -0.653, p > 0.05), and THA-DP-lh (CIMS vs HC: t = -3.345, p < 0.001; CPMS vs HC: t = -2.996, p = 0.003; CIMS vs CPMS: t = -0.366, p > 0.05). No differences were observed in MD between the three groups (p > 0.05). Additionally, both the CIMS and CPMS groups exhibited significant TD increases compared to HC in THA-VPl-rh and THA-VPl-lh, but no differences were found between CIMS and CPMS in any of the thalamic nuclei (Figure 5B): THA-VPl-rh (CIMS vs HC: t = 3.986, p < 0.001; CPMS vs HC: t = 3.918, p < 0.001; CIMS vs CPMS: t = 0.053, p > 0.05), and THA-VPl-lh (CIMS vs HC: t = 3.431, p < 0.001; CPMS vs HC: t = 3.633, p < 0.001; CIMS vs CPMS: t = -0.300, p > 0.05).

**Figure 5.**
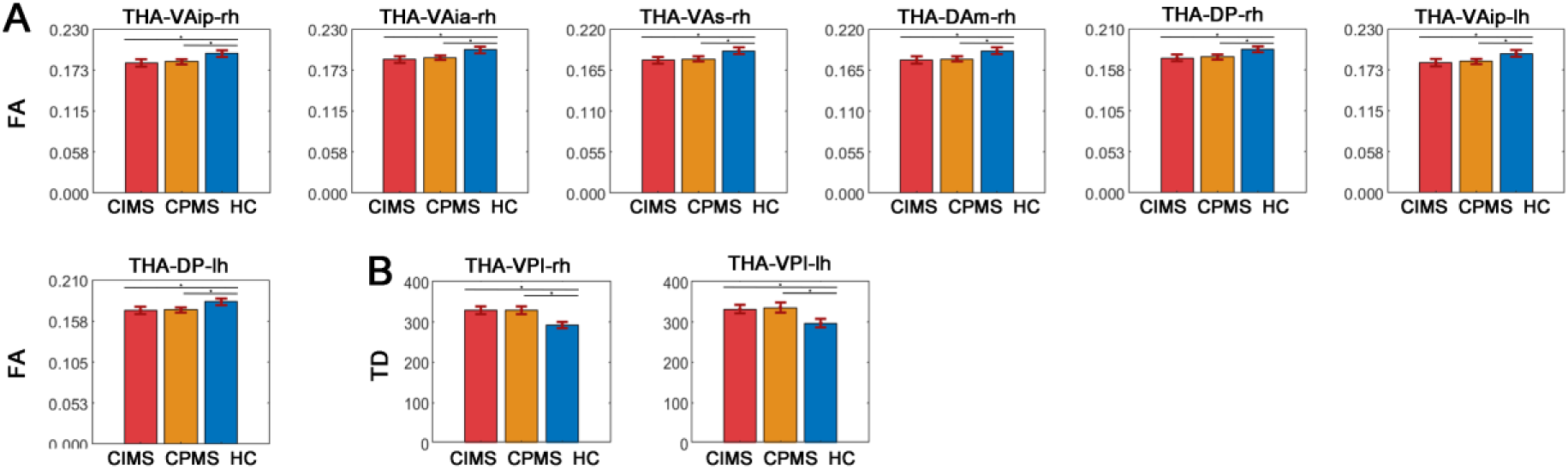
Thalamic diffusion changes in CIMS and CPMS. (5A) Both the CIMS and CPMS groups exhibited significant FA decreases compared to HC in THA-VAip-rh, THA-VAia-rh, THA-VAs-rh, THA-DAm-rh, THA-DP-rh, THA-VAip-lh, and THA-DP-lh, but no differences were found between CIMS and CPMS in any of the thalamic nuclei. **(5B)** Both the CIMS and CPMS groups exhibited significant TD increases compared to HC in THA-VPl-rh and THA-VPl-lh, but no differences were found between CIMS and CPMS in any of the thalamic nuclei.

#### Thalamic controllability metrics changes in CIMS and CPMS

While both the CIMS and CPMS groups exhibited significant increases in average controllability compared to HC in THA-DAm-rh and THA-DAl-rh, the CIMS group showed significantly higher average controllability than CPMS in THA-DAm-rh (Figure 6A): THA-DAm-rh (CIMS vs HC: t = 4.182, p < 0.001; CPMS vs HC: t = 2.615, p = 0.008; CIMS vs CPMS: t = 2.445, p = 0.016), THA-DAl-rh (CIMS vs HC: t = 3.682, p < 0.001; CPMS vs HC: t = 2.903, p = 0.003; CIMS vs CPMS: t = 0.869, p > 0.05). Besides, while both the CIMS and CPMS groups exhibited significant decreases in modal controllability compared to HC in THA-DAm-rh and THA-DAl-rh, the CIMS group showed significantly lower modal controllability than CPMS in THA-DAm-rh (Figure 6B): THA-DAm-rh (CIMS vs HC: t = -4.433, p < 0.001; CPMS vs HC: t = -2.579, p = 0.011; CIMS vs CPMS: t = -2.594, p = 0.009), THA-DAl-rh (CIMS vs HC: t = -3.315, p < 0.001; CPMS vs HC: t = -2.462, p = 0.014; CIMS vs CPMS: t = -0.962, p > 0.05). Additionally, while both the CIMS and CPMS groups exhibited significant decreases in activation energy compared to HC in THA-DAm-rh and THA-DAl-rh, the CIMS group showed significantly lower activation energy than CPMS in THA-DAm-rh (Figure 6C): THA-DAm-rh (CIMS vs HC: t = -4.169, p < 0.001; CPMS vs HC: t = - 2.455, p = 0.015; CIMS vs CPMS: t = -2.546, p = 0.012), THA-DAl-rh (CIMS vs HC: t = -3.736, p < 0.001; CPMS vs HC: t = -3.240, p = 0.002; CIMS vs CPMS: t = -0.549, p > 0.05).

**Figure 6.**
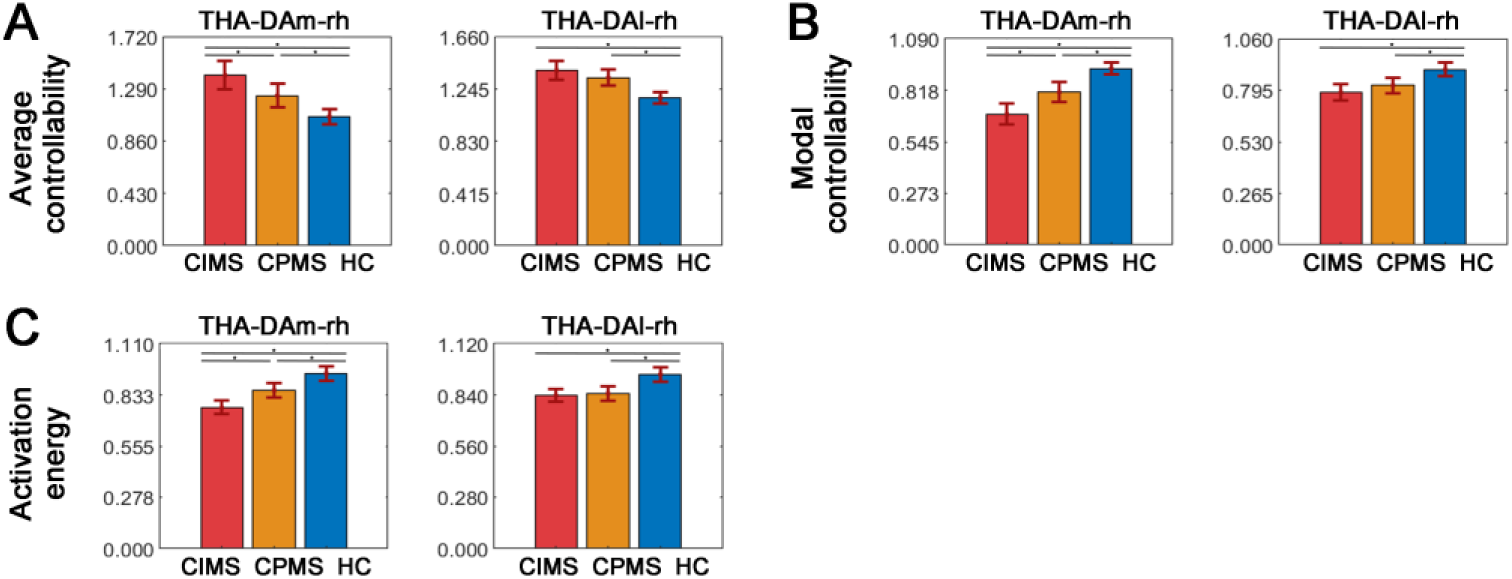
Thalamic controllability changes in CIMS and CPMS. (6A) While both the CIMS and CPMS groups exhibited significant increases in average controllability compared to HC in THA-DAm-rh and THA-DAl-rh, the CIMS group showed significantly higher average controllability than CPMS in THA-DAm-rh. **(6B)** While both the CIMS and CPMS groups exhibited significant decreases in modal controllability compared to HC in THA-DAm-rh and THA-DAl-rh, the CIMS group showed significantly lower modal controllability than CPMS in THA-DAm-rh. **(6C)** While both the CIMS and CPMS groups exhibited significant decreases in activation energy compared to HC in THA-DAm-rh and THA-DAl-rh, the CIMS group showed significantly lower activation energy than CPMS in THA-DAm-rh.

### Associations between changed thalamic imaging metrics and lesion load or cognitive outputs

#### Associations between changed thalamic volume metrics and lesion load or cognitive outputs

Associations between changed thalamic volume metrics and lesion load are shown in Figure 7A. Specifically, significant negative correlations were observed between lesion load and thalamic volume in all thalamic nuclei: THA-VAip-rh (rho = -0.573, *p* < 0.001), THA-VAia-rh (rho = -0.564, *p* < 0.001), THA-VPm-rh (rho = -0.601, *p* < 0.001), THA-VPl-rh (rho = -0.570, *p* < 0.001), THA-VAs-rh (rho = -0.599, *p* < 0.001), THA-DAm-rh (rho = -0.612, *p* < 0.001), THA-DAl-rh (rho = -0.603, *p* < 0.001), THA-DP-rh (rho = -0.523, *p* < 0.001), THA-VAip-lh (rho = - 0.548, *p* < 0.001), THA-VAia-lh (rho = -0.515, *p* < 0.001), THA-VPm-lh (rho = -0.579, *p* < 0.001), THA-VPl-lh (rho = -0.620, *p* < 0.001), THA-VAs-lh (rho = -0.561, *p* < 0.001), THA-DAm-lh (rho = -0.573, *p* < 0.001), THA-DAl-lh (rho = -0.627, *p* < 0.001), and THA-DP-lh (rho = - 0.629, *p* < 0.001).

**Figure 7.**
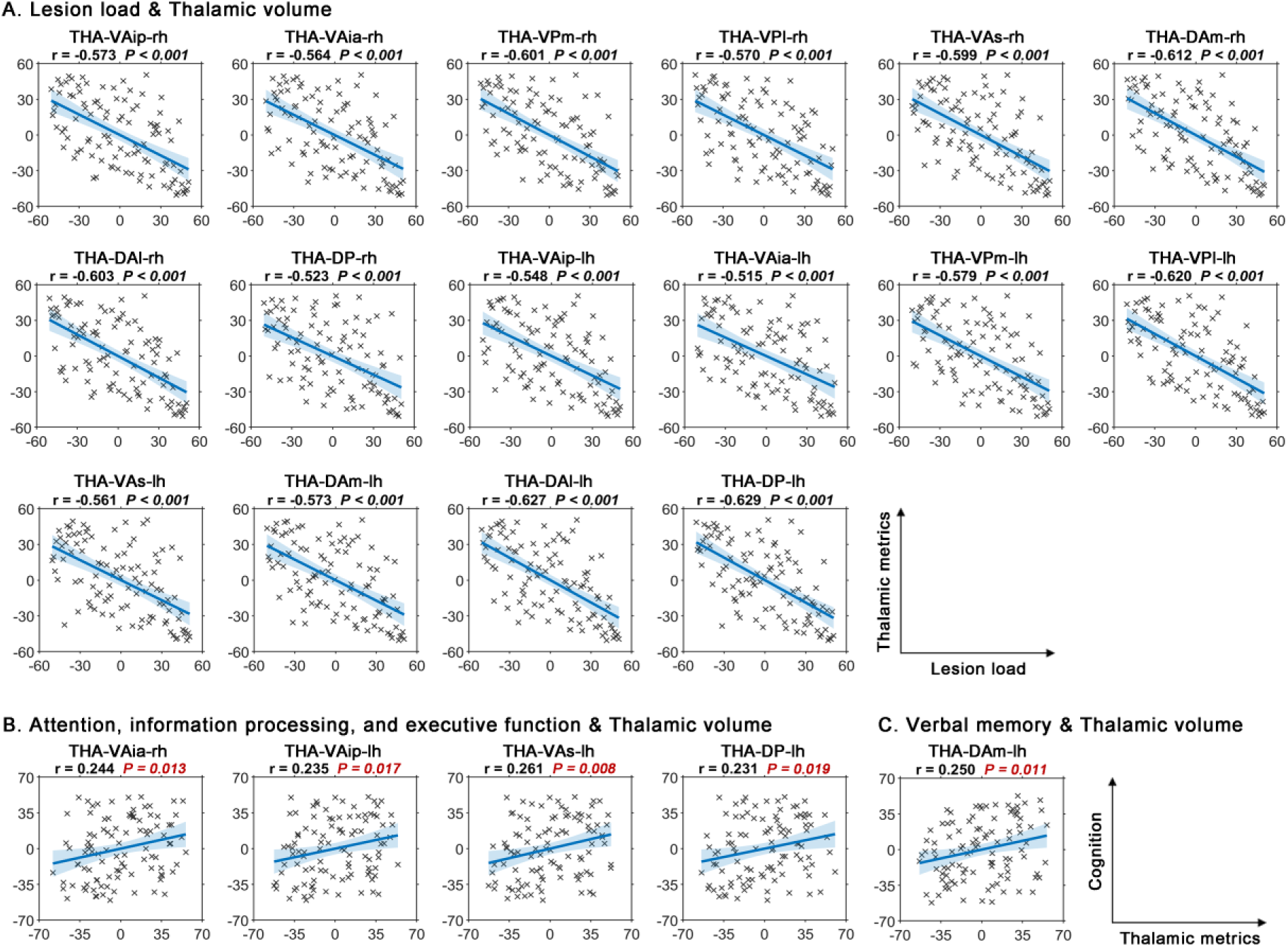
Associations between changed thalamic volume metrics and lesion load or cognitive outputs. (7A) **S**ignificant negative correlations were observed between lesion load and thalamic volume in all thalamic nuclei. **(7B)** Trend-level positive correlations were observed between performance in attention, information processing, and executive function and thalamic volumes in THA-VAia-rh, THA-VAip-lh, THA-VAs-lh, and THA-DP-lh. **(7C)** Trend-level positive correlations were observed between performance in verbal memory and thalamic volumes in THA-DAm-lh.

Associations between changed thalamic volume metrics and cognitive outputs are shown in Figure 7B. Specifically, trend-level positive correlations were observed between performance in attention, information processing, and executive function and thalamic volumes in THA-VAia-rh, THA-VAip-lh, THA-VAs-lh, and THA-DP-lh (*p* < 0.05, uncorrected); as well as between performance in verbal memory and thalamic volumes in THA-DAm-lh (*p* < 0.05, uncorrected); however, these correlations did not reach significance after correction for multiple comparisons.

#### Associations between changed thalamic diffusion metrics and lesion load or cognitive outputs

Associations between changed thalamic diffusion metrics and lesion load are shown in Figure 8. Specifically, significant positive correlation was observed between lesion load and thalamic MD in THA-VAia-lh (rho = 0.411, *p* < 0.001; Figure 8A). Besides, significant negative correlation was observed between lesion load and thalamic TD in THA-DAm-lh (rho = -0.332, *p* < 0.001; Figure 8B). Additionally, trend-level positive correlations were observed between lesion load and thalamic TD in THA-VPl-rh and THA-VPl-lh (*p* < 0.05, uncorrected; Figure 8B); as well as trend-level negative correlations between lesion load and thalamic FA in THA-VAia-rh, THA-DP-rh, THA-VAip-lh, THA-VPm-lh, THA-VPl-lh, THA-DAm-lh, THA-DAl-lh, and THA-DP-lh (*p* < 0.05, uncorrected; Figure 8C); however, these correlations did not reach significance after correction for multiple comparisons.

**Figure 8.**
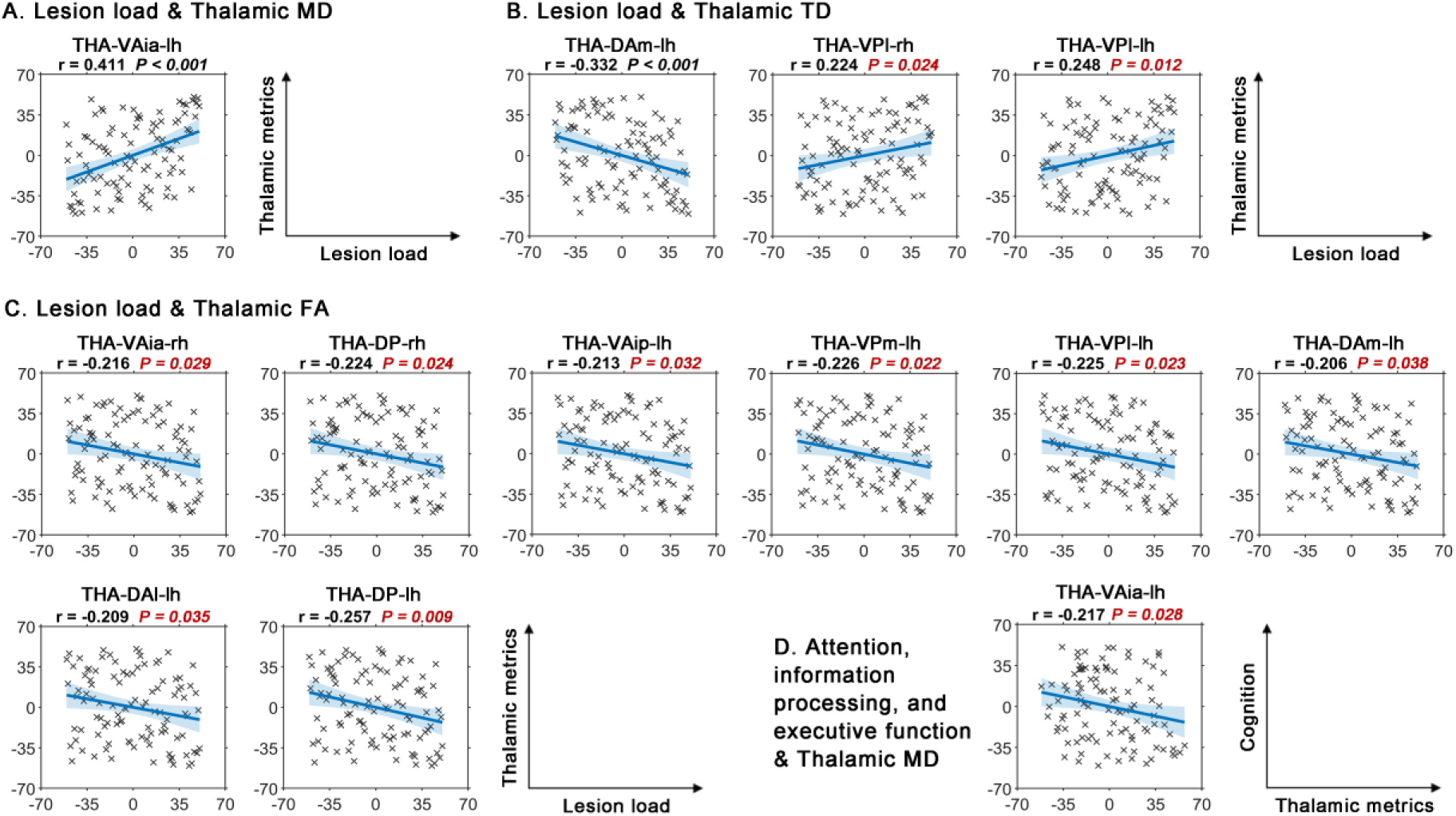
Associations between changed thalamic diffusion metrics and lesion load or cognitive outputs. (8A) **S**ignificant positive correlation was observed between lesion load and thalamic MD in THA-VAia-lh. **(8B)** Significant negative correlation was observed between lesion load and thalamic TD in THA-DAm-lh. Additionally, trend-level positive correlations were observed between lesion load and thalamic TD in THA-VPl-rh and THA-VPl-lh. **(8C)** Trend-level negative correlations between lesion load and thalamic FA in THA-VAia-rh, THA-DP-rh, THA-VAip-lh, THA-VPm-lh, THA-VPl-lh, THA-DAm-lh, THA-DAl-lh, and THA-DP-lh. **(8D)** Trend-level negative correlation was observed between performance in attention, information processing and executive function and thalamic MD in THA-VAia-lh.

Associations between changed thalamic diffusion metrics and cognitive outputs are shown in Figure 8D. Specifically, trend-level negative correlation was observed between performance in attention, information processing and executive function and thalamic MD in THA-VAia-lh (*p* < 0.05, uncorrected); however, the correlation did not reach significance after correction for multiple comparisons.

#### Associations between changed thalamic controllability metrics and lesion load or cognitive outputs

No significant or trend-level associations were observed between changed thalamic controllability metrics and lesion load (p > 0.05).

Associations between changed thalamic controllability metrics and cognitive outputs are shown in Figure 9. Specifically, trend-level negative correlation was observed between performance in verbal fluency and thalamic average controllability in THA-DAm-rh (*p* < 0.05, uncorrected). Besides, trend-level positive correlations were observed between performance in verbal fluency and thalamic modal controllability in THA-DAm-rh (*p* < 0.05, uncorrected) as well as thalamic activation energy in THA-DAm-rh (*p* < 0.05, uncorrected). However, these correlations did not reach significance after correction for multiple comparisons.

**Figure 9.**
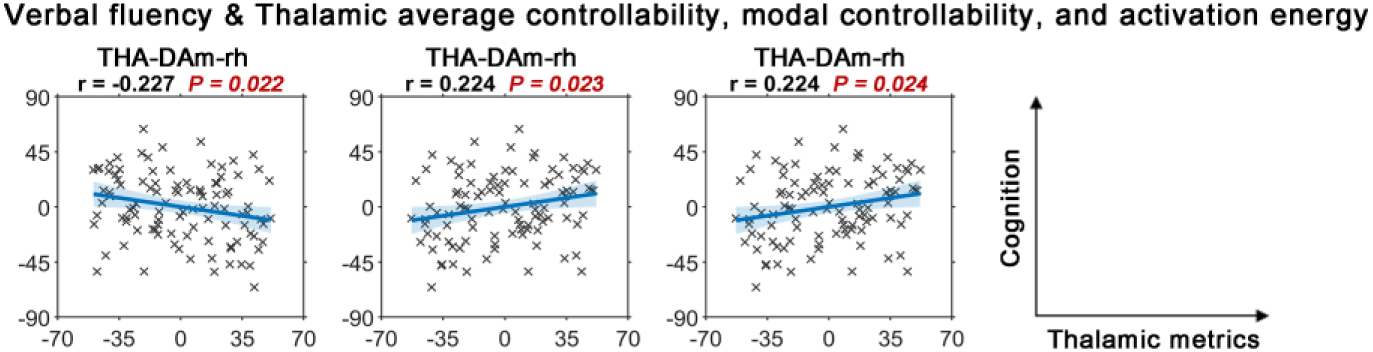
Associations between changed thalamic controllability metrics and lesion load or cognitive outputs. No significant or trend-level associations were observed between changed thalamic controllability metrics and lesion load. Trend-level negative correlation was observed between performance in verbal fluency and thalamic average controllability in THA-DAm-rh. Besides, trend-level positive correlations were observed between performance in verbal fluency and thalamic modal controllability in THA-DAm-rh as well as thalamic activation energy in THA-DAm-rh.

### Associations among changed thalamic imaging metrics

Significant correlations were found between volume and diffusion metrics (Figure 10). Specifically, significant positive correlations were found between thalamic volume and thalamic FA in THA-VPm-rh (rho = 0.372, *p* < 0.001), THA-DAm-rh (rho = 0.296, *p* = 0.003), THA-DP-rh (rho = 0.554, *p* < 0.001), THA-VPm-lh (rho = 0.390, *p* < 0.001), and THA-DP-lh (rho = 0.460, *p* < 0.001) (Figure 10A). Besides, significant negative correlations were found between thalamic volume and thalamic MD in all thalamic nuclei: THA-VAip-rh (rho = -0.481, *p* < 0.001), THA-VAia-rh (rho = -0.508, *p* < 0.001), THA-VPm-rh (rho = -0.358, *p* < 0.001), THA-VPl-rh (rho = -0.405, *p* < 0.001), THA-VAs-rh (rho = -0.473, *p* < 0.001), THA-DAm-rh (rho = - 0.460, *p* < 0.001), THA-DAl-rh (rho = -0.416, *p* < 0.001), THA-DP-rh (rho = -0.302, *p* = 0.003), THA-VAip-lh (rho = -0.483, *p* < 0.001), THA-VAia-lh (rho = -0.501, *p* < 0.001), THA-VPm-lh (rho = -0.341, *p* < 0.001), THA-VPl-lh (rho = -0.482, *p* < 0.001), THA-VAs-lh (rho = -0.431, *p* <, THA-DAm-lh (rho = -0.471, *p* < 0.001), THA-DAl-lh (rho = -0.478, *p* < 0.001), and THA-DP-lh (rho = -0.440, *p* < 0.001) (Figure 10B). Additionally, significant correlations were found between thalamic volume and thalamic TD in almost all thalamic nuclei: THA-VAip-rh (rho = - 0.331, *p* < 0.001), THA-VPm-rh (rho = -0.295, *p* = 0.003), THA-VPl-rh (rho = -0.287, *p* = 0.003), THA-DAm-rh (rho = 0.497, *p* < 0.001), THA-DAl-rh (rho = -0.330, *p* < 0.001), THA-DP-rh (rho = 0.345, *p* < 0.001), THA-VAip-lh (rho = -0.383, *p* < 0.001), THA-VPm-lh (rho = -0.301, *p* =, THA-VPl-lh (rho = -0.399, *p* < 0.001), THA-DAm-lh (rho = 0.361, *p* < 0.001), and THA-DAl-lh (rho = -0.432, *p* < 0.001) (Figure 10C).

**Figure 10.**
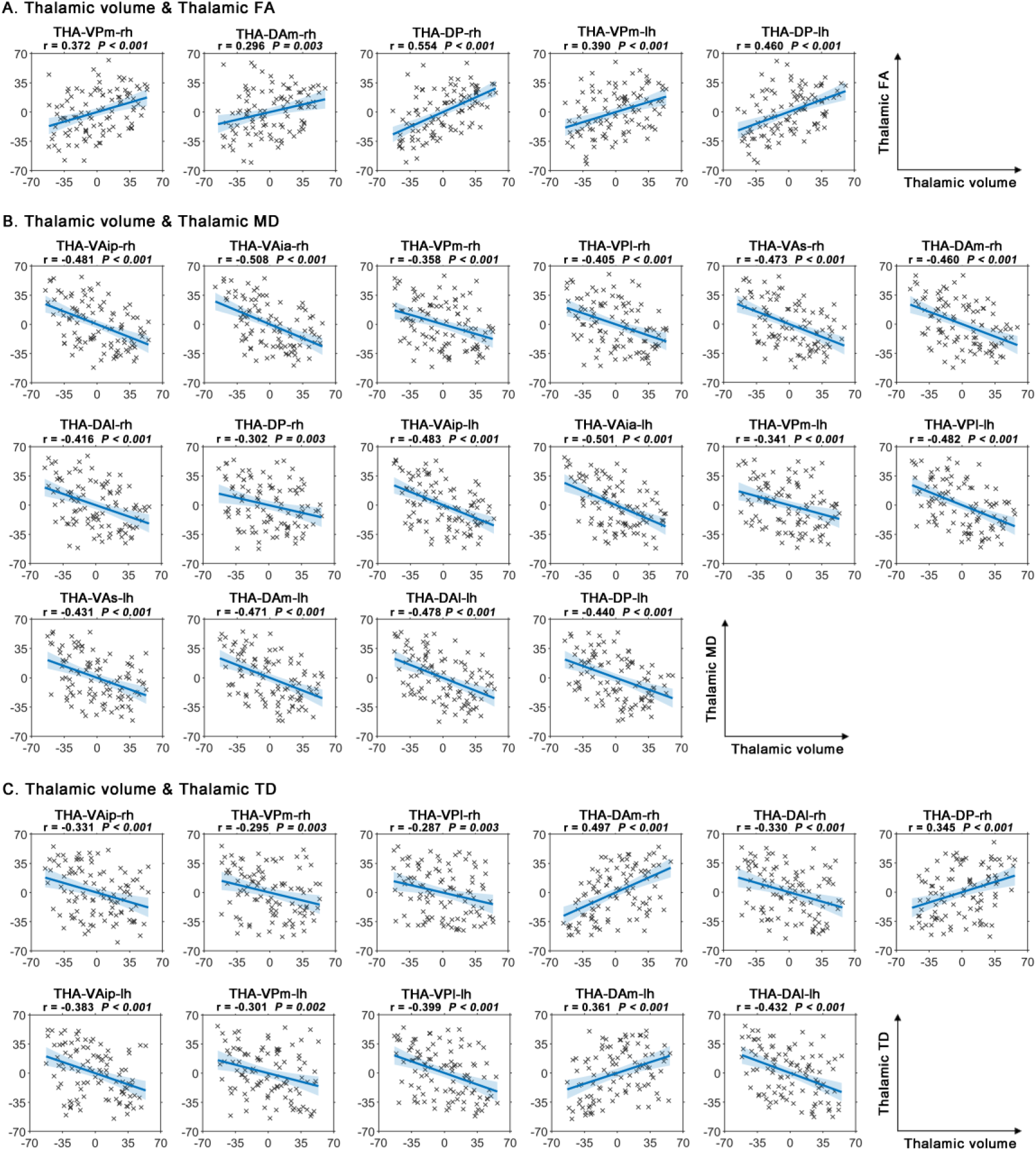
Associations among changed thalamic imaging metrics. (10A) **S**ignificant positive correlations were found between thalamic volume and thalamic FA in THA-VPm-rh, THA-DAm-rh, THA-DP-rh, THA-VPm-lh, and THA-DP-lh. **(10B)** Significant negative correlations were found between thalamic volume and thalamic MD in all thalamic nuclei. **(10C)** Significant correlations were found between thalamic volume and thalamic TD in almost all thalamic nuclei: THA-VAip-rh, THA-VPm-rh, THA-VPl-rh, THA-DAm-rh, THA-DAl-rh, THA-DP-rh, THA-VAip-lh , THA-VPm-lh, THA-VPl-lh, THA-DAm-lh, and THA-DAl-lh. No significant correlations were found between controllability metrics and either volume or diffusion metrics.

No significant correlations were found between controllability metrics and either volume or diffusion metrics (*p* > 0.05).

### Covariance between thalamic imaging metrics and cognitive performance

#### Verbal memory

Controllability metrics exhibited the highest covariance with performance in verbal memory (Figure 11A, Supplementary Table 1). Specifically, the strongest associations were observed with the triple-modality model combining controllability, diffusion, and volume metrics (R = 0.896). Among dual-modality models, the combination of controllability and diffusion metrics yielded the highest association (R = 0.880), followed by controllability and volume (R = 0.788), and diffusion and volume (R = 0.720). For single-modality models, controllability metrics showed the strongest associations (R = 0.705), followed by diffusion (R = 0.674), and volume metrics (R = 0.485).

**Figure 11.**
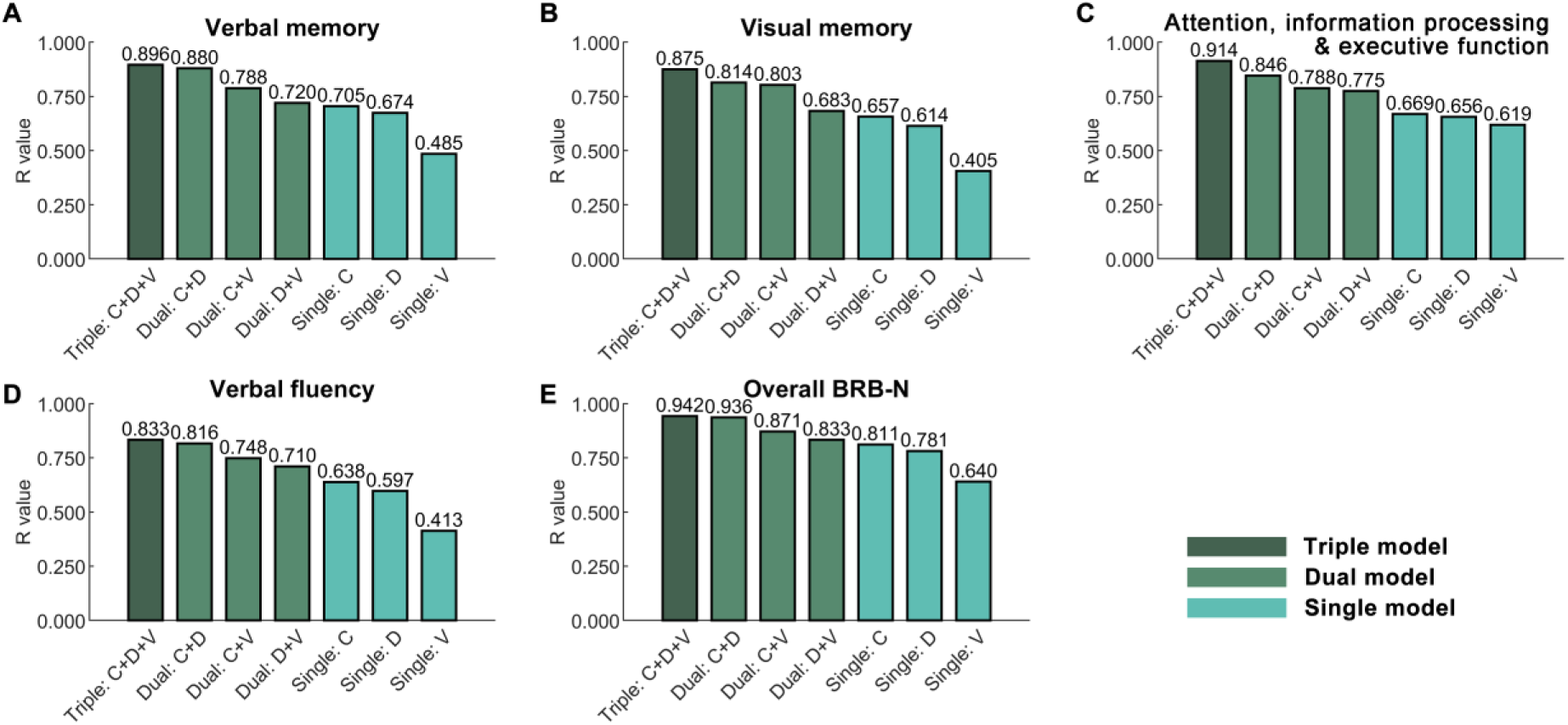
Covariance between thalamic imaging metrics and cognitive performance. (11A) Controllability metrics exhibited the highest covariance with performance in verbal memory. **(11B)** Controllability metrics exhibited the highest covariance with performance in visual memory performance. **(11C)** Controllability metrics exhibited the highest covariance with performance in attention, information processing, and executive function. **(11D)** Controllability metrics exhibited the highest covariance with performance in verbal fluency. **(11E)** Controllability metrics exhibited the highest covariance with performance in overall BRB-N.

#### Visual memory

Controllability metrics exhibited the highest covariance with performance in visual memory performance (Figure 11B, Supplementary Table 2). Specifically, the strongest associations were observed with the triple-modality model combining controllability, diffusion, and volume metrics (R = 0.875). Among dual-modality models, the combination of controllability and diffusion metrics yielded the highest association (R = 0.814), followed by controllability and volume (R = 0.803), and diffusion and volume (R = 0.683). For single-modality models, controllability metrics showed the strongest associations (R = 0.657), followed by diffusion (R = 0.614), and volume metrics (R = 0.405).

#### Attention, information processing, and executive function

Controllability metrics exhibited the highest covariance with performance in attention, information processing, and executive function (Figure 11C, Supplementary Table 3). Specifically, the strongest associations were observed with the triple-modality model combining controllability, diffusion, and volume metrics (R = 0.914). Among dual-modality models, the combination of controllability and diffusion metrics yielded the highest association (R = 0.846), followed by controllability and volume (R = 0.788), and diffusion and volume (R = 0.775). For single-modality models, controllability metrics showed the strongest associations (R = 0.669), followed by diffusion (R = 0.656), and volume metrics (R = 0.619).

#### Verbal fluency

Controllability metrics exhibited the highest covariance with performance in verbal fluency (Figure 11D, Supplementary Table 4). Specifically, the strongest associations were observed with the triple-modality model combining controllability, diffusion, and volume metrics (R = 0.833). Among dual-modality models, the combination of controllability and diffusion metrics yielded the highest association (R = 0.816), followed by controllability and volume (R = 0.748), and diffusion and volume (R = 0.710). For single-modality models, controllability metrics showed the strongest associations (R = 0.638), followed by diffusion (R = 0.597), and volume metrics (R = 0.413).

#### Overall BRB-N

Controllability metrics exhibited the highest covariance with performance in overall BRB-N (Figure 11E, Supplementary Table 5). Specifically, the strongest associations were observed with the triple-modality model combining controllability, diffusion, and volume metrics (R = 0.942). Among dual-modality models, the combination of controllability and diffusion metrics yielded the highest association (R = 0.936), followed by controllability and volume (R = 0.871), and diffusion and volume (R = 0.833). For single-modality models, controllability metrics showed the strongest associations (R = 0.811), followed by diffusion (R = 0.781), and volume metrics (R = 0.640).

## Discussion

In this study, we examined the functional controllability of specific thalamic nuclei in individuals with MS. We investigated whether alterations in controllability metrics were associated with changes in grey and white matter structure within these nuclei. We found that individuals with MS exhibited widespread thalamic grey matter atrophy and white matter FA decreases across all nuclei, whereas changes in functional controllability were selectively observed in the dorsal anterior thalamic nuclei compared to healthy controls. Moreover, functional controllability in the medial dorsal anterior nuclei differed between cognitively impaired and cognitively preserved MS groups. Notably, individual variability in controllability metrics was independent of thalamic grey matter volume, white matter integrity, and lesion load. Crucially, combining thalamic functional controllability with structural metrics yielded a stronger association with cognitive performance in MS than either modality alone.

Our findings revealed widespread grey matter atrophy across all thalamic nuclei in MS. We also observed associations between thalamic volume reductions and poorer cognitive performance, particularly in attention, executive function, and verbal memory, aligning with prior studies. Structural MRI research has consistently linked thalamic grey matter atrophy to cognitive impairment in MS (Amin & Ontaneda, 2021; Bergsland et al., 2016, 2021; Bisecco et al., 2021; Conway et al., 2021; Houtchens et al., 2007; Kletenik et al., 2019; Lorefice et al., 2020; Rocca et al., 2010; Rojas et al., 2018; Sandroff et al., 2022; Schoonheim et al., 2015; Štecková et al., 2014; Valdés Cabrera et al., 2022). A recent meta-analysis reviewed structural MRI-derived morphological measures across the brain associated with cognitive impairment in MS, and concluded that thalamic volume shows the strongest associations with various cognitive domains, including information processing speed, visuospatial learning and memory, verbal learning and memory, and executive function (Mirmosayyeb et al., 2024). However, it is worth noting that the associations we observed did not survive Bonferroni correction (p < 0.05, uncorrected), likely due to the multiple comparisons across multiple subnuclei rather than the thalamus as a whole, which may have reduced statistical power and resulted in trend-level findings.

We observed widespread reductions in thalamic white matter FA across all nuclei in MS. Besides, significant increases in MD in THA-VAia-lh as well as increases in TD in THA-VPl-rh and THA-VPl-lh as well as decreases in THA-DAm-lh were also observed in MS. Notably, increased MD in THA-VAia-lh showed a trend-level correlation with reduced performance in attention and executive function. These findings are consistent with previous diffusion-weighted imaging studies, which have commonly reported reduced FA and increased MD in the thalamus in MS. For example, MS-related decreases in thalamic FA has been associated with deficits in global cognitive performance, particularly in attention, information processing speed, and verbal memory (Schoonheim et al., 2014), while MS-related increases in thalamic MD has been linked to poorer performance in verbal learning and visuospatial memory (Bergsland et al., 2018). In our study, only thalamic MD—but not FA or TD—showed associations with cognitive performance. This aligns with prior work suggesting that MD may be a more sensitive marker of cognitive decline in MS than FA (Benedict et al., 2013; Schoonheim et al., 2015).

Unlike the widespread structural changes observed across all thalamic nuclei, functional controllability alterations were concentrated in the dorsal anterior nuclei, where MS showed increased average controllability, decreased modal controllability, and decreased activation energy compared to HC. Notably, although structural measures did not differentiate between cognitively impaired and preserved individuals with MS, functional controllability metrics did: cognitively impaired individuals exhibited more pronounced changes in functional controllability in thalamic medial dorsal anterior nuclei, as demonstrated by significant higher average controllability and lower modal controllability and lower activation energy when compared to the cognitively preserved individuals. Our previous study (Yang et al., 2025) demonstrated that thalamic functional controllability can differentiate cognitive subgroups in MS, the current findings advance this by identifying the medial dorsal anterior nuclei as the most affected subregion. The medial dorsal thalamic nuclei has been reported to drive input directly from various regions of the prefrontal cortex and maintain strong connections with limbic system structures (Mitchell & Chakraborty, 2013; Parnaudeau et al., 2018). These anatomical features underpin their critical role in a range of cognitive functions, including memory, decision-making, and executive processes, as demonstrated by comparative data across multiple species (Mitchell & Chakraborty, 2013; Parnaudeau et al., 2018). Moreover, the medial dorsal thalamus has been implicated in several neurological and psychiatric disorders, such as stroke, dementia, schizophrenia, major depressive disorder, Parkinson’s disease, and Alzheimer’s disease (Mitchell & Chakraborty, 2013; Parnaudeau et al., 2018). Our findings extend this literature by identifying its involvement in cognitive impairment associated with MS.

Notably, we observed trend-level associations between increased average controllability, decreased modal controllability, and decreased activation energy in the medial dorsal thalamic nuclei and poorer verbal fluency performance—a subdomain of language processing—in individuals with MS. In contrast, a recent study using the same methodology in a healthy cohort (Yang et al., 2025. bioRxiv) found that higher average controllability, lower modal controllability, and lower activation energy in the dorsal anterior thalamic nuclei were associated with better language processing. In line with this, another study demonstrated that acute low-frequency stimulation of the dorsal subthalamic nucleus improved verbal fluency in individuals with Parkinson’s disease (Lee et al., 2021), further supporting the role of dorsal thalamic nuclei in modulating language function. Therefore, we hypothesise that individuals with MS and impaired verbal fluency may engage in a compensatory adaptation, whereby they attempt to enhance average controllability while reducing modal controllability and activation energy in the dorsal thalamic nuclei in an effort to partially offset deficits in language processing. Future longitudinal studies are needed to validate this hypothesis and examine how functional controllability in the dorsal thalamic nuclei evolves alongside cognitive decline related to language processing.

Our data showed strong correlations between thalamic grey matter volume and white matter integrity measures, suggesting interdependence between structural changes. However, functional controllability metrics were not associated with any structural metric, including grey matter volume, white matter FA, MD, or TD. This suggests that thalamic functional controllability alterations are independent of structural changes in MS. Previous studies have reported that thalamic volume, MD, and functional connectivity each independently predicted cognitive performance in MS using linear regression models (Schoonheim et al., 2015), Extending this, our study directly tested the independence of structural and functional metrics by assessing their correlations at the subnuclei level.

Furthermore, we advanced beyond traditional functional connectivity analyses confined to a single cognitive state by examining dynamic functional control across multiple cognitive states (Deng et al., 2022; Gu et al., 2015). This approach offers novel insights into how individual thalamic nuclei contribute to cognitive flexibility from a dynamic control perspective—capturing aspects of thalamic function that are not reflected in structural properties alone. More importantly, while structural grey and white matter alterations correlated with lesion load, functional controllability did not, further indicating that functional network changes could result from secondary neurodegeneration rather than local inflammation. This independence highlights the potential of functional controllability to capture mechanisms underlying cognitive impairment that are not detectable via conventional structural or lesion-based measures.

Our results demonstrate that thalamic functional controllability metrics exhibited the strongest associations with cognitive performance in MS across all assessed domains— verbal memory, visual memory, attention and executive function, and verbal fluency— compared to structural metrics alone. Notably, combining functional controllability with grey and white matter structural metrics yielded even stronger associations with cognitive performance than any single modality. This finding is consistent with our previous study using the same methodology in a healthy cohort (Yang et al., 2025, bioRxiv. Chapter 5), supporting the view that thalamic functional controllability provides unique and complementary information beyond thalamic volume and microstructural integrity in both healthy and MS populations. The thalamus has long been recognised as a central hub that plays a critical role in various cognitive processes (reviewed in (Shine et al., 2023; Wolff & Vann, 2019)), particularly in MS (reviewed in (Amin & Ontaneda, 2021; DeLuca et al., 2015; Kipp et al., 2015)). A recent work has highlighted the thalamus’s involvement in facilitating cognitive flexibility by regulating transitions between cognitive states in the healthy brain (Rikhye et al., 2018). By applying network controllability analysis, our study offers novel evidence for the importance of thalamic nuclei in MS-related cognitive impairment from the perspective of dynamic control during cognitive processing. Collectively, these findings underscore the potential of functional controllability as a powerful tool for advancing our understanding of the complex role of the thalamus in cognitive function.

This study has some limitations. First, our analyses were restricted to structural and resting-state functional MRI data. While informative, task-based fMRI may offer additional perspectives on cognitive dysfunction in MS and should be explored in future studies.

Second, we utilised a single thalamic parcellation approach, which may not capture the full complexity of thalamic organisation. Employing alternative or multiple parcellation frameworks in future work could help validate the present findings and enhance their generalisability. Third, we focused on linear associations between imaging metrics and cognitive outcomes. Future research should consider applying more sophisticated analytical models to investigate potential nonlinear or interactive effects, which may better reflect the complexity of brain–cognition relationships in MS. Finally, the cross-sectional nature of this study precludes any conclusions about temporal dynamics or causality. Longitudinal designs will be essential to determine how changes in thalamic functional controllability relate to the trajectory of cognitive decline in MS.

## Conclusion

This study provides novel evidence that functional controllability within the thalamus, particularly the medial dorsal anterior nuclei, contributes to cognitive impairment in multiple sclerosis (MS). While widespread grey matter atrophy and white matter microstructural changes were observed across all thalamic nuclei, functional controllability alterations were more spatially selective in the medial dorsal anterior nuclei and most pronounced in cognitively impaired individuals. Importantly, functional controllability metrics were independent of structural measures and lesion load, highlighting their distinct contribution to brain dysfunction in MS. Moreover, combining thalamic functional controllability with structural metrics yielded stronger associations with cognitive performance than either modality alone. By applying a network control framework, our findings offer a dynamic perspective on how changes in thalamic nuclei may relate to cognitive deficits in MS. Overall, this study highlights the potential of integrating structural and functional controllability measures to improve our understanding of the role of thalamic nuclei in MS and cognitive impairment.

## Data and Code Availability

The data and code that support the results of this study are available from the corresponding author upon reasonable request and with permission.

## Author Contributions

Y.L., N.T.B., and N.M. contributed to the conception and design of the study. I.L., V.T., and N.M. contributed to the acquisition of data. Y.Y., J.H. and R.K. contributed to analysis or interpretation of data. Y.Y., A.W., V.T., Y.L., N.T.B., and N.M. contributed to drafting/revision of the manuscript.

## Acknowledgements

This work was funded by research grants from the MS Society UK and Medical Research Council UK (MR/X005267/1). We extend our gratitude to all our colleagues from the MISC and SPiN labs for their kind support during this study.

## Supplementary Material

Supplementary materials are available online.

## Abbreviations

BRB-N: Brief Repeatable Battery of Neuropsychological Tests
CCA: canonical correlation analysis
CIMS: cognitively impaired multiple sclerosis
CPMS: cognitively preserved multiple sclerosis
CSF: cerebrospinal fluid
dMRI: diffusion MRI
EDSS: Expanded Disability Status Scale
EPI: echo planar imaging
FA: fractional anisotropy
FAST: FSL’s Automated Segmentation Tool
FWHM: full width at half maximum
GM: grey matter
HC: healthy controls
MD: mean diffusivity
MS: multiple sclerosis
MSFC: Multiple Sclerosis Functional Composite
ROI: region of interest
rs-fMRI: resting-state functional MRI
SD: standard deviation
T1w: T1-weighted structural MRI
TD: track density
WM: white matter.

