## Supplementary Materials for "Thalamic Nuclei Functional Controllability Accounts for Cognitive Impairment in Multiple Sclerosis Over and Above Structural Damage"

**Supplementary Table 1. Covariance between thalamic imaging metrics and performance in verbal memory.**

| Modality | Thalamic imaging metrics | R |
|---|---|---|
| Triple-modality model | Controllability + Diffusion + Volume | 0.896 |
| Dual-modality model | Controllability + Diffusion | 0.880 |
| | Controllability + Volume | 0.788 |
| | Diffusion + Volume | 0.720 |
| Single-modality model | Controllability | 0.705 |
| | Diffusion | 0.674 |
| | Volume | 0.485 |

R values are generated using sparse canonical correlation analysis.

**Supplementary Table 2. Covariance between thalamic imaging metrics and performance in visual memory.**

| Modality | Thalamic imaging metrics | R |
|---|---|---|
| Triple-modality model | Controllability + Diffusion + Volume | 0.875 |
| Dual-modality model | Controllability + Diffusion | 0.814 |
| | Controllability + Volume | 0.803 |
| | Diffusion + Volume | 0.683 |
| Single-modality model | Controllability | 0.657 |
| | Diffusion | 0.614 |
| | Volume | 0.405 |

R values are generated using sparse canonical correlation analysis.

**Supplementary Table 3. Covariance between thalamic imaging metrics and performance in attention and executive function.**

| Modality | Thalamic imaging metrics | R |
|---|---|---|
| Triple-modality model | Controllability + Diffusion + Volume | 0.914 |
| Dual-modality model | Controllability + Diffusion | 0.846 |
| | Controllability + Volume | 0.788 |
| | Diffusion + Volume | 0.775 |
| Single-modality model | Controllability | 0.669 |
| | Diffusion | 0.656 |
| | Volume | 0.619 |

R values are generated using sparse canonical correlation analysis.

**Supplementary Table 4. Covariance between thalamic imaging metrics and performance in verbal fluency**

| Modality | Thalamic imaging metrics | R |
|---|---|---|
| Triple-modality model | Controllability + Diffusion + Volume | 0.833 |
| Dual-modality model | Controllability + Diffusion | 0.816 |
| | Controllability + Volume | 0.748 |
| | Diffusion + Volume | 0.710 |
| Single-modality model | Controllability | 0.638 |
| | Diffusion | 0.597 |
| | Volume | 0.413 |

R values are generated using sparse canonical correlation analysis.

**Supplementary Table 5. Covariance between thalamic imaging metrics and performance in overall BRB-N.**

| Modality | Thalamic imaging metrics | R |
|---|---|---|
| Triple-modality model | Controllability + Diffusion + Volume | 0.942 |
| Dual-modality model | Controllability + Diffusion | 0.936 |
| | Controllability + Volume | 0.871 |
| | Diffusion + Volume | 0.833 |
| Single-modality model | Controllability | 0.811 |
| | Diffusion | 0.781 |
| | Volume | 0.640 |

R values are generated using sparse canonical correlation analysis.
